# Population genomics provide insights into the global genetic structure of *Colletotrichum graminicola,* the causal agent of maize anthracnose

**DOI:** 10.1101/2022.05.07.490914

**Authors:** Flávia Rogério, Riccardo Baroncelli, Francisco Borja Cuevas-Fernández, Sioly Becerra, JoAnne Crouch, Wagner Bettiol, M. Andrea Azcárate-Peril, Martha Malapi-Wight, Veronique Ortega, Javier Betran, Albert Tenuta, José S. Dambolena, Paul D. Esker, Pedro Revilla, Tamra A. Jackson-Ziems, Jürg Hiltbrunner, Gary Munkvold, Ivica Buhiniček, José L. Vicente-Villardón, Serenella A. Sukno, Michael R. Thon

## Abstract

**Background:** *Colletotrichum graminicola*, the causal agent of maize anthracnose, is an important crop disease worldwide. Understanding the genetic diversity and mechanisms underlying genetic variation in pathogen populations is crucial to the development of effective control strategies. The genus *Colletotrichum* is largely recognized as asexual, but several species have been reported to have a sexual cycle. Here, we employed a population genomics approach to investigate the genetic diversity and reproductive biology of *C. graminicola* isolates infecting maize. We sequenced 108 isolates of *C. graminicola* collected in 14 countries using restriction site-associated DNA sequencing (RAD-Seq) and whole-genome sequencing (WGS).

**Results:** Clustering analyses based on single-nucleotide polymorphisms showed populational differentiation at a global scale, with three genetic groups delimited by continental origin, compatible with short-dispersal of the pathogen, and geographic subdivision. Distinct levels of genetic diversity were observed between these clades, suggesting different evolutionary histories. Intra and inter-continental migration was predicted between Europe and South America, likely associated with the movement of contaminated germplasm. Low clonality and evidence of genetic recombination were detected from the analysis of linkage disequilibrium and the pairwise homoplasy index (PHI) test for clonality. We show evidence that even if rare (possibly due to losses of sex and meiosis-associated genes) *C. graminicola* can undergo sexual recombination based on lab assays and genomic analyses.

**Conclusions:** Our results support hypotheses of intra and intercontinental pathogen migration and genetic recombination with great impact on *C. graminicola* population structure.

## Introduction

Crop diseases are one of the biggest concerns in global agriculture responsible for major losses in food production [1,2]. Fungi are primary causal agents of plant diseases and can cause catastrophic damage [3,4]. Disease outbreaks caused by fungal plant pathogens during recent decades have increased and are recognized as emergent threats to food security worldwide [5–8]. Human activity tends to intensify fungal disease dispersal by promoting pathogen movement far from their native location [9,10]. The modification of natural environments caused by modern agriculture provides new opportunities for natural selection [10]. The employ of crops with a limited genetic diversity increases the risks of global disease spread since pathogen genotypes can rapidly spread through genetically uniform host populations [11,12]. Additionally, ongoing climate changes can shift plant-pathogen distributions on a global scale since temperature changes may affect diseases development, hence acting in the spread and survival of pathogens and their establishment in hitherto unsuitable regions [8,13,14].

Maize (*Zea mays*) is the second most extensively cultivated cereal crop globally, with a harvested area of almost 200 million hectares [15]. Together with rice and wheat, maize supplies nearly half daily calories in human food [14,16]. Although many diseases negatively impact maize production, anthracnose is one of the most important diseases, it is so difficult to control that requires to use of integrated practices [17–20]. Anthracnose disease is caused by *Colletotrichum graminicola*, an ascomycete fungus that can infect most plant tissues, though the stalk rot and seedling blight can bring more significant economic damage [21–23]. Taxonomically, *C. graminicola* belongs to the Graminicola species complex, a well-defined monophyletic clade encompassing 16 *Colletotrichum* species pathogenic to the Poaceae family [23–25].

Prior to 1960, maize anthracnose was a minor leaf spot disease in the USA, and the causal agent *C. graminicola* was primarily studied for its taxonomic significance [22,26]. However, this view changed during the following decades when devastating outbreaks caused unexpected losses across north-central and eastern regions of the USA [19,22]. Afterwards, the disease became increasingly common, and in some seasons extremely destructive throughout the eastern USA [27–29]. The occurrence of these outbreaks are thought to be associated with the use of hybrids particularly susceptible to anthracnose, more favorable weather conditions, and the emergence of more aggressive strains of the pathogen with rapid spread through the maize belt [30–32]. Nowadays, maize anthracnose is a well-established endemic disease in the USA, and better managed mainly through the use of resistant cultivars, but it still can cause important damage in some regions of the country [33–37]. Given the history of the disease, it may have the potential to become much more significant in the future.

Outside of the USA*, C. graminicola* is globally distributed and is of major importance in the Americas, and other regions of the world [38]. In North America, Canada reports variable disease incidence, but anthracnose is still the cause of significant concern [36,39,40]. In addition to temperate areas, maize anthracnose is also reported in tropical and subtropical regions, including Brazil and Argentina, where warm and humid conditions contribute to augment the disease [41–48]. Since the first report of anthracnose in Europe, (Italy in 1852) [49], several countries have reported the occurrence of the disease, including Germany, France, and ex Czechoslovakia [50–52], and more recently Croatia, Portugal, Switzerland, Bosnia and Herzegovina [53–57]. Recently, maize anthracnose also was reported in Asia [58], indicating that the pathogen is increasing its geographic range.

Pathogens widely disseminated over continents indicate an intense dispersal of their propagules, either from aerial dispersal, or via human-mediated movement of infected plant material, and/or seeds [11,59–61]. The nature of fungal propagules, i.e., their small and numerous spores, favor long-distance dispersal resulting in the spread of pathogens across and between continents [11]). Although many pathogens are cosmopolitan in distribution, their populations often can differ in some aspects, for instance, virulence, resistance to chemical controls, and genetic diversity [62]. The migration of plant pathogens can have epidemiological and evolutionary implications since the movement of individuals from one population to another may result in gene flow when migrants contribute their genes to the gene pool of another population [63]. *Colletotrichum graminicola* is seed-transmitted, also known as a seed-born pathogen [64–66], therefore, contaminated seeds serve as inoculum sources, and promote pathogen movement. Furthermore, though the transport of seeds and infected plant material, fungi can naturally be spread via the dispersal of both sexual and asexual spores [11,59,67]. In addition to the dispersal process, the reproductive biology of a species is crucial to understanding the evolutionary forces that shape the genetic diversity of pathogen populations over time and space [68]. Asexual reproduction predominates in the most of plant-pathogenic fungi, but many species undergo sexual cycles, and many others employ both reproductive strategies [63]. Sexual reproduction produces genetic recombination by the independent assortment of chromosomes, increasing genotype diversity and generating a random association between alleles at different loci [69,70]. Yet, alternative mechanisms, such as parasexual reproduction, and cryptic sexual cycle can also create genetic recombination in fungal species [71,72]. While sexual reproduction may be costly (in terms of fitness) and disrupt favorable gene combinations, experimental evidence has suggested that sex in fungi may increase the rate of adaptation to new environments [73,74]. Recombination can create particular genotypes through the combination of more fit alleles, which can accelerate the evolution of resistance to multiple fungicides and overtake R genes of resistant cultivars [69,75,76].

The genetic control underlying sexual development in the genus *Colletotrichum* is not well known and differs from the typical bipolar mating system of ascomycetes [77–79]. All species studied so far contain just one of the two mating types alleles typical of ascomycetes (*MAT1-1/2*) [23], which would seemingly limit their ability to mate. However, for at least some *Colletotrichum* species, including *C. graminicola* sexual reproduction has been demonstrated. Although the teleomorph of *C. graminicola* has been characterized under laboratory conditions [78,79], the asexual state is the one commonly found in the field conditions, which plays a significant role in the lifecycle. This species is reported to be a homothallic species [78], i.e., possesses both sets of genes and thus can complete the sexual cycle independently [80]. However, some strains are self-sterile and cross-fertile [79,81], which means that some strains are capable of reproducing sexually when cultured alone, while other isolates require a mating partner to produce sexual offspring.

Genetic recombination is reported as an important factor in the ecology and evolution of many *Colletotrichum* species [82–87]. However, only a few population studies have characterized the genetic diversity of grass-infecting *Colletotrichum* species, such as *C. sublineola* and*, C. cereale* [85–89], and little is known about *C. graminicola* population diversity. Studies using single marker genes and limited sample size showed high genetic variability on *C. graminicola* isolates [87,90]. Therefore, in this study, we investigated the global population structure of *C. graminicola* to get insights into the reproductive biology of this species. Our population genomic analyses using single-nucleotide polymorphisms (SNPs) revealed population subdivision into three genetic clades with isolates grouped according to their country of origin, with distinct levels of genetic diversity between them, suggesting different evolutionary histories. We found intra and intercontinental migration, mainly between Europe and South America, likely associated with the movement of contaminated seeds. We discussed the possible evolutionary origins of these genetic groups and the impact of recombination on the genetic structure as well the potential of maize anthracnose becoming a much more significant disease in the future.

## Materials And Methods

### Sampling and isolation of *C. graminicola*

Fresh collections of *C. graminicola* samples were obtained from symptomatic maize stems provided by an international network of maize researchers and producers, allowed sampling from nine different countries where anthracnose disease is prevalent (Fig. 1). Eleven *C. graminicola* isolates originally collected from five countries (Germany, Japan, Netherlands, Nigeria, and Zimbabwe) [91,92] were obtained from public culture collections (Supplementary Table S1) and also included in the collection. Symptomatic stems were dissected, surface disinfected, and plated in culture medium to fungal isolation, as previously described [21,57]. Fungal structures (hyphal tips, acervuli) emerging from maize tissue were transferred to potato dextrose agar (PDA) medium allowed to grow for 7 days at 23°C then single-spore purified. Monosporic cultures were stored at -80°C from suspension conidia in 7.5% skim milk (Merck, Darmstadt Germany) mixed on silica gel (S/0730/53 Fisher) [93]. Isolates were verified as *C. graminicola* by inspection of colony morphology, spore size and shape, and sequencing of the internal transcribed spacer (*ITS*) region of rDNA using the primers ITS4 and ITS5 [94], following PCR amplification conditions as described by Rech et al. [85]. The amplified ITS fragments were purified using NucleoSpin gel and PCR Clean-up kits (Macherey Nagel, Düren, Germany) and sequenced using Sanger technology by University of Salamanca (Spain). Specie identity was confirmed by BLAST searches performed against in GenBank (NCBI). All materials used in this study were used in accordance with the rights and obligations of the recipient as described in the International Treaty on Plant Genetic Resources for Food and Agriculture adopted by the FAO on November 3, 2001.

**Figure 1.**
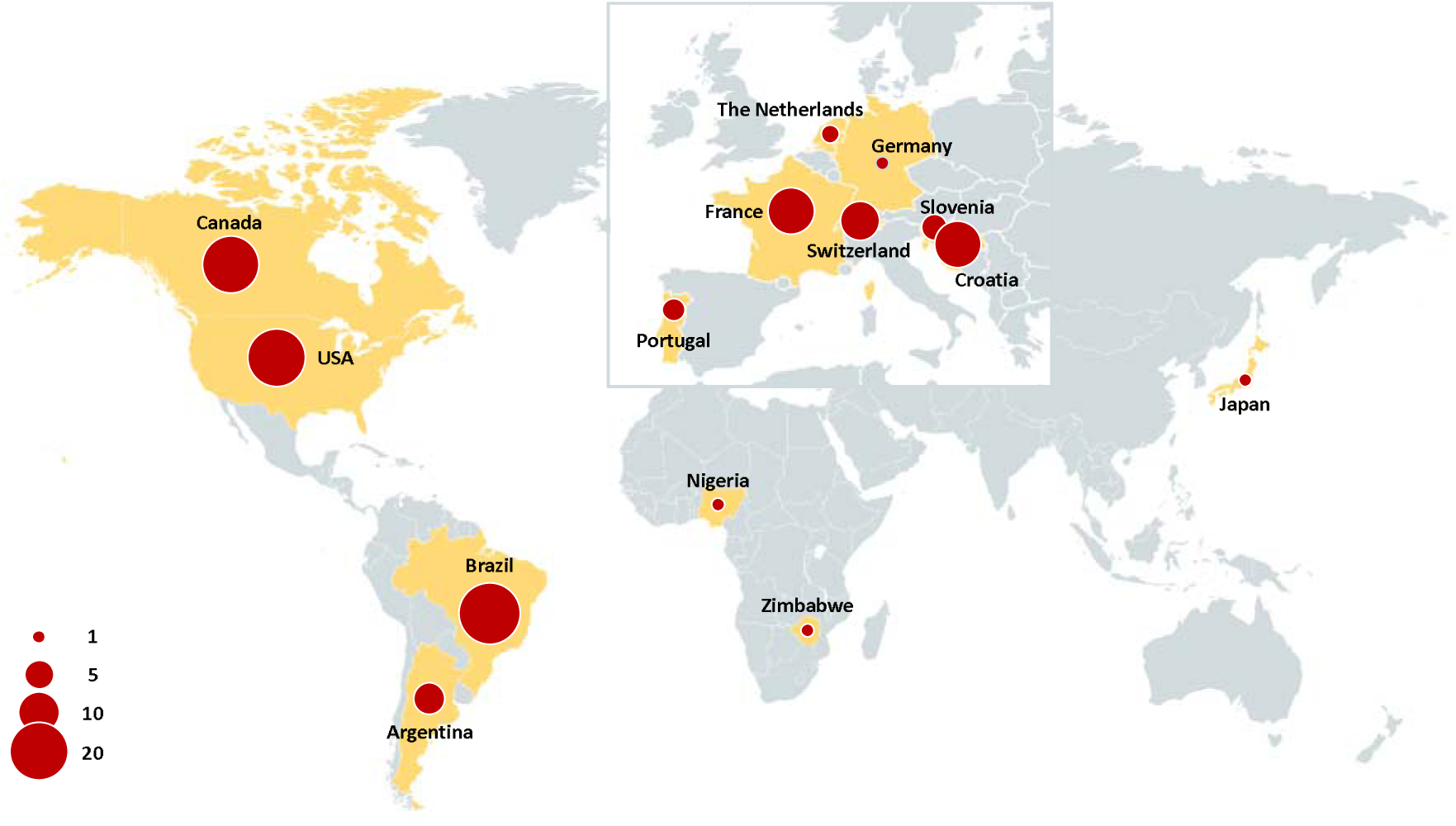
Worldwide *Colletotrichum graminicola* sampling.

### DNA extraction, library preparation, and sequencing

Mycelia from monosporic cultures were incubated on an orbital shaker in potato dextrose broth (PDB) (Difco, Becton, Dickinson and Company, Franklin Lakes, USA) for three days at 25°C and 150 rpm under continuous light [54]. Genomic DNA was extracted using DNeasy Plant Mini Kits (Qiagen Inc., Valencia, California, USA) and quantified using a Qubit 3.0 fluorimeter (Invitrogen, Waltham, USA). DNA for *C. graminicola* isolates used for whole genome sequencing (WGS) was extracted using the cetyltrimethylammonium-bromide (CTAB) protocol adapted from Baek and Kernerley [95].

New DNA sequence datasets were generated from 101 *C. graminicola* isolates using either RAD-seq or WGS; sequences for an additional seven isolates were also included in the analyses [92] (Supplementary Table S1, S2). A total of 88 isolates were genotyped with RAD-seq by Floragenex Inc. (Beaverton, Oregon, USA). Briefly, RAD-seq libraries with sample-specific barcode sequences were constructed from DNA digested with restriction enzyme PstI, then single-end sequenced (1 x 100 bp) on one lane using TrueSeq PE150 kit (Illumina, Inc., San Diego, CA) on an Illumina HiSeq 2000 instrument. WGS data was obtained from eight *C. graminicola* isolates (CBS252.59, F-64330-2, F-64330-7, F-40330-1, SW-8046-2, CRO-I-35, CR-10370-50, SW-8046-2) from libraries prepared using the Nextera XT DNA kit (Illumina, Inc, San Diego, CA, and sequenced on Illumina HiSeq4000 (2×150 bp). WGS data was obtained from six *C. graminicola* isolates (ARG-2700-2, CA-CHAT-1, CA-N8H, CRO-I-41, P-7565-072-1, SW-8046-1) from paired-end libraries prepared using the Illumina TruSeq DNA Nano kit and sequenced on an Illumina MiSeq (2×300 bp). Sequence quality was checked on FASTQC v0.11.7 (Babraham Bioinformatics) and low-quality reads and adaptors were trimmed with trimmomatic v0.3.8 [96]. *De novo* assemblies were performed with SPADES v3.11.1 [97]. Low coverage scaffolds were identified and removed; scaffolds from the mitochondrial genome and rRNA clusters were identified and masked. The assembly completeness was assessed based on the sordariomyceta_odb9 lineage dataset using BUSCO v.3 [98].

### Read mapping and SNP calling

The RAD-Seq reads were inspected with FASTQC v0.11.7 (Babraham Bioinformatics, Cambridge, UK) to confirm quality and were demultiplexed and quality filtered with the process_radtags program in STACKS v.2.10 [99]. The demultiplexed reads were trimmed using trimmomatic v.0.3.8 [96] using the parameters (’LEADING:15 TRAILING:15 SLIDINGWINDOW:4:15 MINLEN:36’) and aligned to the *C. graminicola* reference genome M1.001 (NCBI accession number SAMN02953757) using BWA v.0.7.8 [100]. Genetic variants were identified with freebayes V.0.9.21 [102]. The WGS reads were quality trimmed with trimmomatic v.0.3.8 and aligned to the reference genome with speedseq V.0.1.2 [101], which uses BWA-MEM for read mapping and freebayes for variant identification. The vcf files were merged and filtered using vcftools [103] to remove nonbiallelic SNPs do not present in all isolates, sites with quality scores lower than 30, and minor allele frequency (MAF) lower than 3%.

### Population structure and genetic diversity

We defined a population as a group of individuals of the same species that occupy that share the same area and colonize the same type of host simultaneously, then sharing a common evolutionary history [63]. Analysis of molecular variance (AMOVA) was performed with the hypothesis of no genetic differentiation using the package poppr v2.8.6 [106]. Weir and Cockerham’s genetic differentiation index (F_ST_) was calculated with the R package hierfstat using the WC84 method [107]. We used three clustering methods to examine the C. *graminicola* population structure. First, we used a principal component analysis (PCA), with *a priori* geographical knowledge from sampling (by continent). This analysis was conducted in the R environment with the package adegenet v2.0.1 [108] using the dudi.pca function. Second, we performed a discriminate analysis of principal components (DAPC) with R package adegenet v2.0.1 [108]. The find.cluster function was used to identify the most likely grouping in the data based on the Bayesian information criterion (BIC) calculated for 1 to 10 clusters. Third, to confirm the pattern of population subdivision inferred by the PCA and DAPC we used the fast maximum-likelihood genetic clustering method snapclust [109]. The function snapclust.choosek was applied as a prior step to determine the optimal number of clusters (*K*), based on the Bayesian information criterion (BIC). Additionally, we implemented a neighbor network using the software splitstree V4 [110] to visualize phylogenetic signals, indicated by the presence of reticulations in the network, from a distance matrix with default parameters.

The number of multilocus genotypes (MLGs) was calculated using the R package poppr v2.8.6 [106]. We used the MGL.FILTER function to determine the true number of MLGs from a genetic distance obtained by DISS.DIST function using the threshold determined in the cutoff_predictor tool. To visualize relationships among genotypes a minimum spanning network (MSN) was generated using the *msn* function in the R package poppr. Genetic diversity was estimated by comparison of the number of SNP variations by counting the number of segregating sites between individuals, using the find.variations tool in geneious [111].

### Linkage disequilibrium

The coefficient of linkage disequilibrium (*r*^2^) [113] was calculated using vcftools [103]. The rate of decay in linkage disequilibrium with physical distance was calculated for all pairs of SNPs less than 100 kb apart using the option --hap-r2 and excluding sites with minor allele frequencies below 5%. We also used the pairwise homoplasy index (PHI) test implemented in splitstree V4 [110] to test the null hypothesis of clonality.

### Investigation of putative sex-related genes

To identify orthologues of sex-related genes in the *C. graminicola* genomes, BLASTp searches (e-value cutoff 1e-3) were performed using for the query a set of 96 gene with a role in sexual reproduction in fungal model systems [114,115]. All nucleotide sequences were checked manually to confirm no interruption in the translation frame. For this analysis, we used the 12 *C. graminicola* assemblies generated in this study (see Supplementary Table S2 for the summary assemblies statistics), plus seven *C. graminicola* genome assemblies used by Rech et al. [92] and genome assemblies from nine other fungal species: *Colletotrichum fiorinae* ([116]; Genome accession: JARH00000000.1), *Colletotrichum fructicola* ([117]; Genome accession: ANPB00000000.1), *Colletotrichum gloeosporioides* ([118]; Genome accession: AMYD00000000.1), *Colletotrichum higginsianum* ([119]; Genome accession: LTAN00000000.1), *Colletotrichum nymphaeae* ([120]; Genome accession: JEMN00000000.1), *Colletotrichum salicis* ([120]; Genome accession: JFFI00000000.1), *Colletotrichum simmondsii* ([120]; Genome accession: JFBX00000000.1), *Colletotrichum sublineola* ([121]; Genome accession: JMSE00000000.1), *Colletotrichum orbiculare* ([117]; Genome accession: AMCV00000000.1), *Verticillium dahliae* ([122]; Genome accession: ABJE00000000.1),

### Pathogenicity assays

Pathogenicity characterization was performed by *in vivo* leaf blight assays using a derivative of the susceptible maize inbred line Mo940 [123]. Plants were cultured in a greenhouse for two weeks (V3 developmental stage), and inoculation via *C. graminicola* spore suspensions were performed according to Vargas et al. [124]. Virulence was quantified by measuring the area of necrotic lesions on leaves 4 days after inoculation. Symptomatic leaves were excised from the plants, scanned with a flat-bed scanner, and quantified using the imaging processing software Paint.NET v.4.2.16 (dotPDN LCC) [54], with three biological replicates per isolate. For the comparison of differences in virulence among isolates, Analysis of Variance (ANOVA) with Bonferroni and Sidak corrections for pairwise comparisons was performed. Seventeen isolates were assessed, divided into batches because we could not manage the whole set simultaneously in the laboratory. Given the possibility of a batch effect, one of the strains (M1.001) was analyzed in all the batches to estimate the possible systematic differences among them. Once all the isolates were analyzed, we compared the isolate repeated in the different batches using ANOVA and obtained highly significant differences (*p* < 0.0001), indicating a clear batch effect. Therefore, to eliminate the systematic differences among batches, we standardized each strain using the mean and the standard deviation of the control of its batch.

## Results

### Population genetic analyses

A total of 5811 biallelic variants were detected among 108 *C. graminicola* isolates collected from 14 countries, which were used for population genetic analyses (Fig. 1; Supplementary Table S1). Analysis of SNP density showed an average of 1.4 SNPs per 10 kb across contigs. Information about reads and coverage per individual are listed in the supplementary table S1.

Clustering methods revealed population subdivision, supporting the existence of three genetic groups, with isolates grouped by their continent of origin corresponding to the isolates from South America, Europe, and North America, henceforth referred to as Brazilian (BR), European (EU), and North American (NAM) clades (Fig. 2). The PCA and DAPC grouped individuals according to their geographic origin (Fig. 2a, Supplementary Fig. S1). Nevertheless, the isolates LAR318, CBS113173, and MAFF511343, from Zimbabwe, Nigeria, and Japan respectively, obtained from culture collections did not group with the main three genetic clusters. We do not discard the possibility of their geographic origins may not be correctly reported in the collections. The SNAPCLUST function based on the BIC criterion supported the clustering pattern by showing the optimum value of *K* = 3 clusters (Supplementary Fig. S2). The neighbor-net network generated by SPLITSTREE identified three groups in the data (Fig. 3). Clusters in the network were connected by extensive reticulated branches (looping) and long branches among samples, indicating a large evolutionary distance separating individuals. AMOVA performed between *C. graminicola* clusters showed that 65.5% of the total genetic variance was within clusters and 34.5% was among them. Pairwise F_ST_ values between clusters were significant and revealed strong differentiation, ranging from 0.26 to 0.42 (Supplementary Table S3).

**Figure 2.**
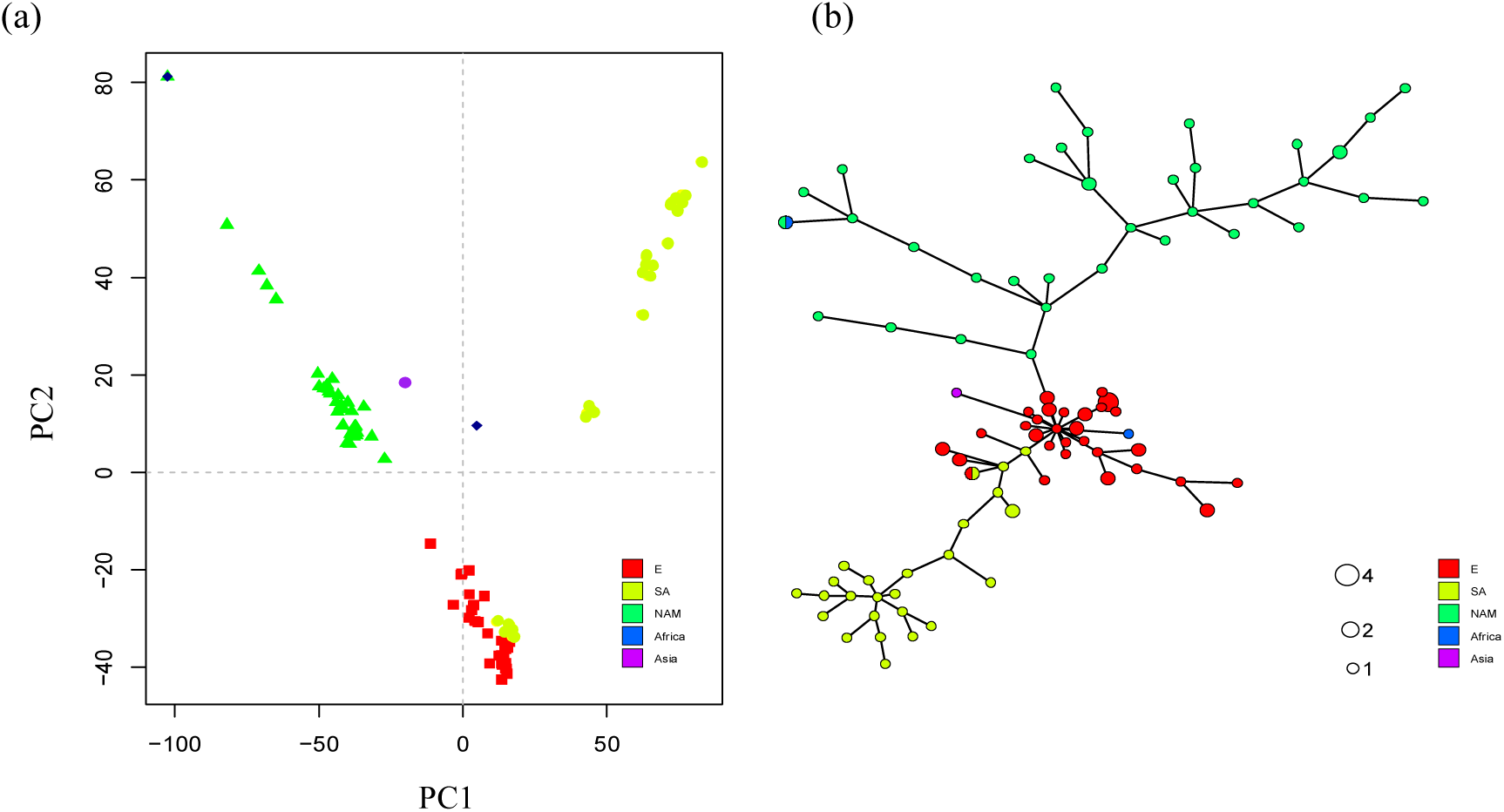
Population subdivision of *Colletotrichum graminicola*. (A) Principal components analysis (PCA). Isolates are colored according to their continent of origin. (B) Minimum spanning network (MSN) showing the multilocus genotypes (MLGs) based on Nei’s genetic distance. The distance between nodes represents the genetic distance between MLGs.

**Figure 3.**
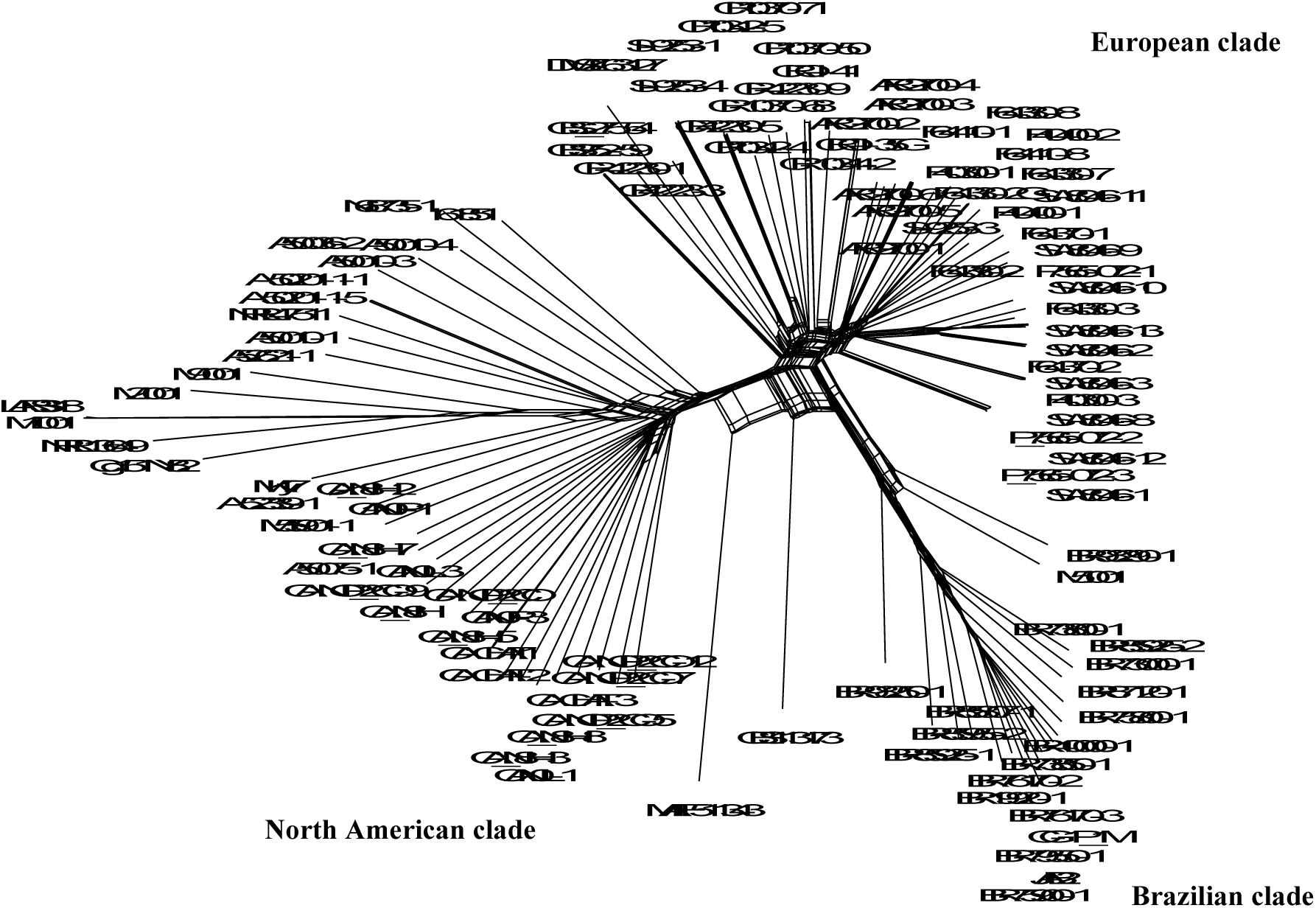
Neighbor-Net network showing relationships between *Colletotrichum graminicola* isolates.

**Figure 4.**
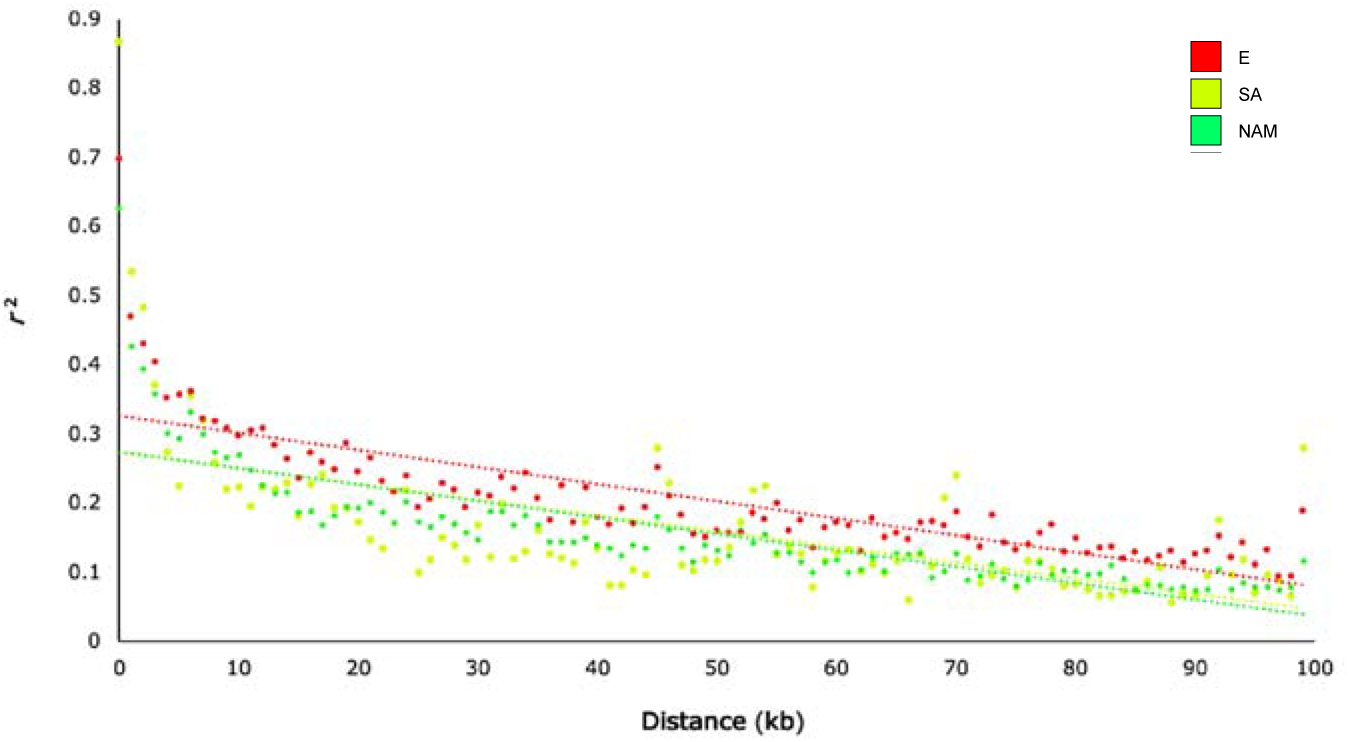
Linkage disequilibrium (LD) decay plots of three genetic populations of *Colletotrichum graminicola* (European, Brazilian, and North America). LD was calculated for all pairs of SNPs less than 100 kb distance apart.

Among the 108 isolates, we detected 76 MLGs unique and 15 MLGs shared between at least two isolates. Most MLGs included isolates from the EU clade (13 MLGs in the total) and shared between isolates from the same clade, and country (Supplementary Table S4). However, one MLG encompassed isolates from France and Switzerland (F-40300-3 and SW-8046-3), and another with isolates from Argentina and Croatia (ARG-2700-2 and CRO-I-35). Two isolates from culture collections (M1.001 and LAR318) from USA and Nigeria were also shared. Such genotypes shared were almost identical genetically, with few SNPs between samples (Supplementary Table S4). The minimum spanning network (MSN) showed a similar relationship among the aforementioned genotypes. The European cluster displayed an inner position with MLGs closer genetically, while Brazilian and North American clusters were placed on the edges of the network, likely due to their separate evolutionary history.

To estimate genetic diversity, we computed the number of segregating sites within genetic groups. Genetic diversity was twice as high in NAM compared to BR and EU clades (Table 1). Based on all the SNPs from the full dataset, NAM showed almost all sites segregating when considering all isolates belonging to this clade. In contrast, isolates from the EU clade showed fewer segregating SNPs and consequently lower genetic diversity.

**Table 1.** Single nucleotide polymorphism (SNP) variation within *Colletotrichum graminicola* clusters.

| Cluster | SNPs within clusters |
| --- | --- |
| BR | 2484 |
| EU | 2077 |
| NAM | 5264 |

\*Standard deviation in parentheses.

### Footprints of recombination

We assessed the genomic impact of recombination by analyzing patterns of linkage disequilibrium. For this, we computed LD on the genetic clusters separately to avoid combining evolutionarily distinct populations, which can lead to an overestimate of the linkage equilibrium [69]. For all genetic clusters, the LD decayed quickly with physical distance, reaching 50% of its maximum value in less than 1kb, suggesting a random association of alleles at different loci, likely caused by recombination events. We also used the pairwise homoplasy index (PHI) test implemented in SPLITSTREE to test the null hypothesis of no recombination, which rejected the null hypothesis for all three genetic populations (*p* < 0.001).

To test the hypothesis that *C. graminicola* sex and meiosis-associated genes are being maintained, we explore the presence of 96 genes previously identified as having roles in sexual reproduction in Ascomycetes (Supplementary Table S5). We observed the presence of 94 genes out of the 96 sex-related genes in three *C. graminicola* genomes (M1.001, LARS318 and CBS113173), while all the other strains were missing three or more genes. A summary of the 15 genes showing the highest variability in presence/absence in the *C. graminicola* genomes is shown in Table 2.

**Table 2.**
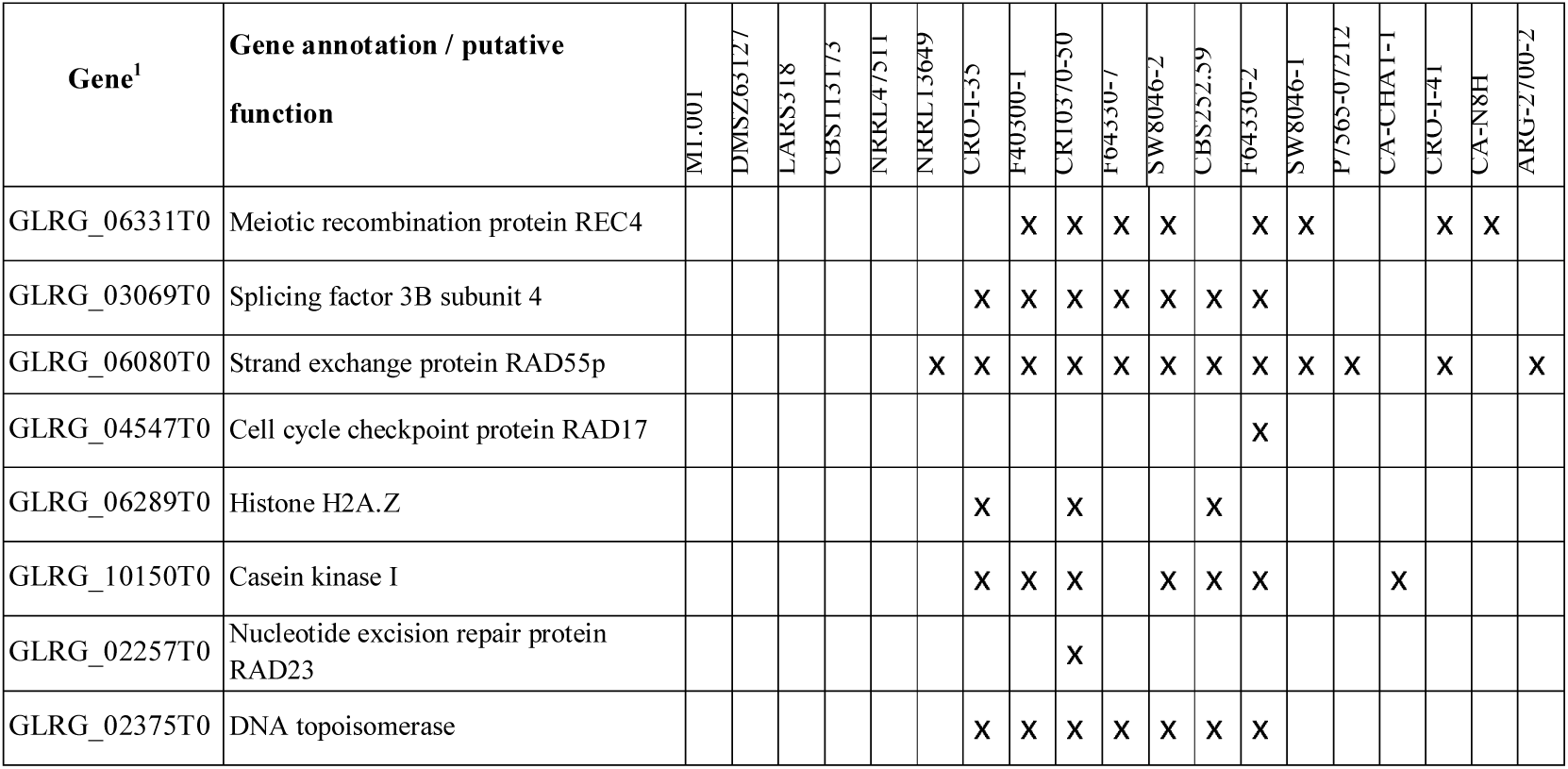
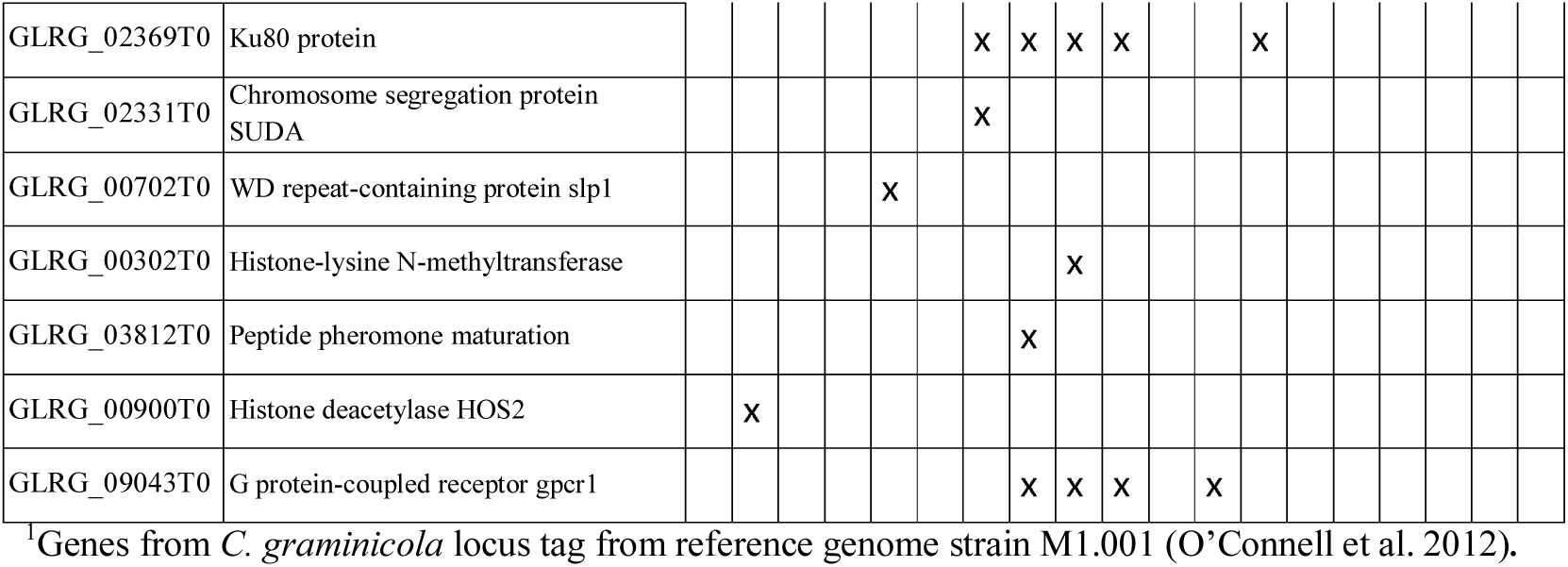
*Colletotrichum graminicola* orthologs of *Fusarium graminearum, Neurospora crassa, Podospora anserina, Saccharomyces cerevisiae*, and *Schizosaccharomyces pombe* sex-related genes. Fifteen genes showing the highest variability in terms of present/absence in the *C. graminicola* genomes. Genes absent are marked with an x.

### Pathogenicity

Seventeen isolates representative of the subclades identified in the phylogeny inference and showing diversity of colony morphology and pigmentation were chosen for pathogenic characterization (Supplementary Fig. S3). Considering the continental origin, the following isolates were chosen, America: M1.001, A-52339-1, ARG-2700-5, BR-85925-1, CA-NOP-2CO, CA-CHAT-1, and JAB2; Africa: CBS113173, LARS318; Europe: CBS252.59, CR-42230-1, CRO-I-41, F-64330-2, P-7565-072-3, SI-9253-1, and SW-8046-2; Asia: MAFF511343. All isolates, except two - CBS252.59 and MAFF511343, were pathogenic on the maize cultivar evaluated producing necrotic lesions at the inoculation points by 4 days post inoculation (dpi) (Fig. 5). Isolates CBS252.59 and MAFF511343 also failed to produce necrotic lesions at 6 dpi (not shown). A high variation in virulence among the isolates was found compared to the strain M1.001. We compared all the strains to the virulence of M1.001 using a classical analysis of variance *p* < 0.0001). Isolates ARG-2700-5, P-7565-072-3, SI-9253-1, CA-CHAT-1, and CR-42230-1 had significantly higher virulence levels, while isolate JAB2 was significantly less virulent. Population structure was not correlated with virulence.

**Figure 5.**
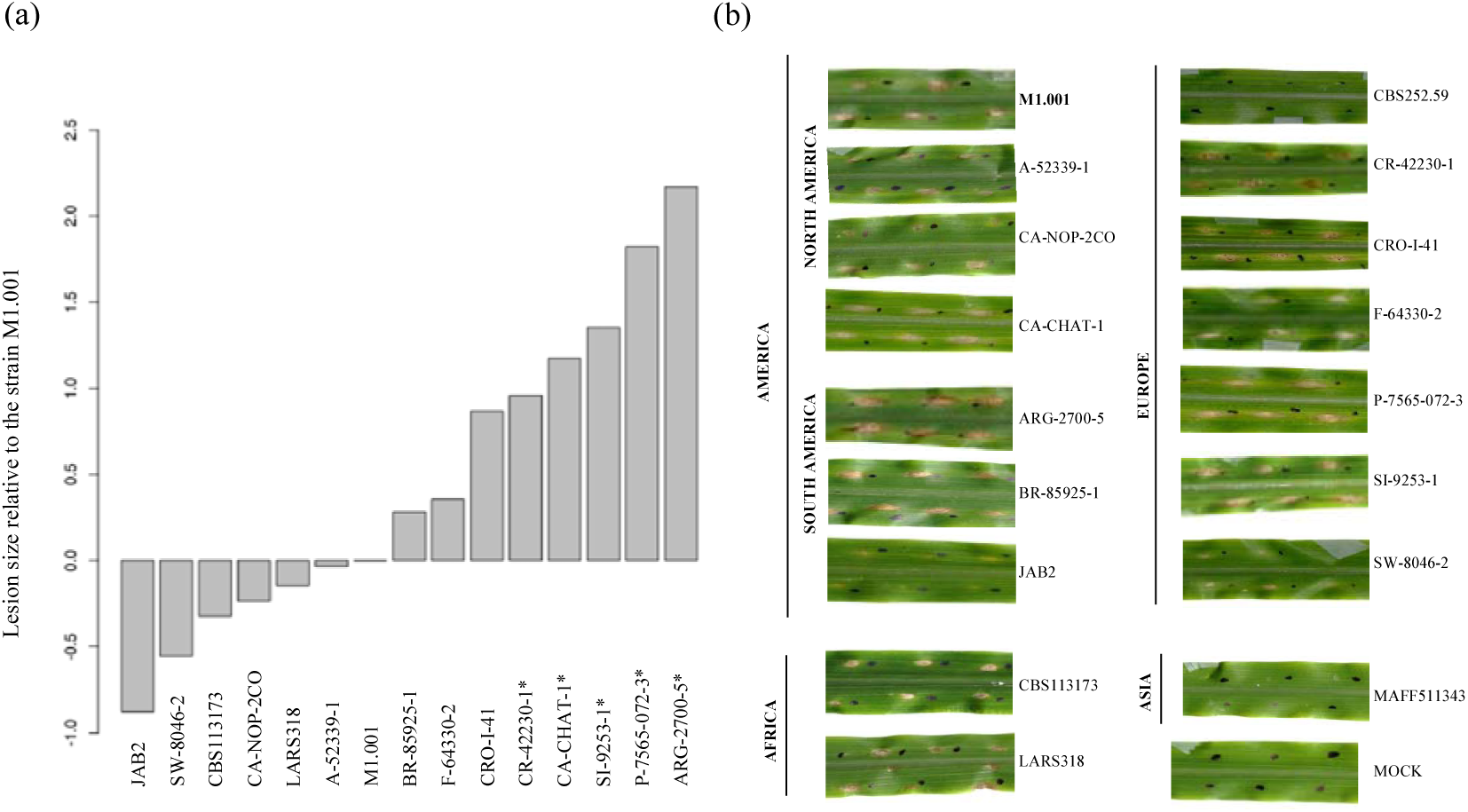
Pathogenic characterization of *Colletotrichum graminicola* isolates. (a) Graphic representation of pairwise comparisons of 14 isolates in order of virulence (relative to strain M1.001 used as control strain) in ascending order. The isolates CBS252.59 and MAFF511343 were non-pathogenic to the maize cultivar evaluated and were not included in the figure. Isolates marked with an asterisk are significantly different from isolate M1.001. (b) Necrotic lesions on maize leaves at 4 days after inoculation with spore suspensions of the isolates.

## Discussion

We employed a population genomics approach to understand the genetic diversity of *C. graminicola* isolates infecting maize sampled from fourteen countries. Using SNP markers generated through RAD-seq and WGS, we investigated the distribution of genetic variation considering our samples as a global population. We found populational differentiation at a worldwide scale, with three genetic groups delimited by continental origin, compatible with short dispersal of the pathogen, and geographic subdivision. However, we observed migration intra and intercontinental migration, likely associated with the movement of contaminated seeds. The genetic clusters, corresponding to Brazilian, European, and North American clades, showed distinct levels of genetic diversity that could reflect different evolutionary histories. Evidence of genetic recombination was detected from analysis of linkage disequilibrium and the PHI test for clonality. Investigation of sex-related genes revealed gene losses in most isolates. We show evidence that even if rare (possibly due to losses of sex and meiosis-associated genes) *C. graminicola* can undergo sexual recombination based on lab assays and genomic analyses. We present the possible evolutionary origins of these genetic groups and the impact of recombination on genetic structure and the potential of maize anthracnose becoming a much more significant disease in the future.

Clustering analyses based on single-nucleotide polymorphisms showed strong population differentiation, grouping of the isolates according to their country of origin. The European clade showed less genetic variation, a higher number of repeated MLGs, and lower pairwise differentiation among clusters, suggesting a larger clonal component as compared to the other clades. Interestingly, all genotypes shared were almost identical genetically, suggesting that gene flow can be dependent on the clonal isolates generated by asexual reproduction. Most MLGs were shared between isolates from the same cluster and country, however, one MLG encompassed isolates from France and Switzerland (more than 1000 km apart) and another one with isolates from Argentina and Croatia. In our cluster analyses, the European clade encompassed all isolates from Argentina. Together, these results indicate intra and intercontinental migration of the pathogen. We argue a likely European origin of the isolates from Argentina due to the movement of maize germplasm into South America. It is common for maize breeders from North America and Europe to use South America (including Argentina) fields for advancing breeding programs during the northern hemisphere winter. As Argentina is a maize exporter, breeding programs can represent a possible source of pathogen importation and consequently a source of pathogen introductions into new areas, for instance in Europe or elsewhere. This result is particularly important from the epidemiological point of view, with direct implications on disease management since migration can cause the movement of more virulent and/or fungicide resistant genotypes.

The predominant form of dissemination of *C. graminicola* is thought to be by contaminated seeds [64–66]. Fungi can naturally be spread via the dispersal of both sexual and asexual spores, caused mainly by the transport of infected plant material [11,59,67]. Dispersal of asexual conidia of *C. graminicola* is strongly dependent on water splash and blowing raindrops, responsible for short-distance dispersal [22]. Although airborne dispersal is not typically documented, ascospore dispersal is associated with long-distance dissemination since such spores can be ejected up and dispersed by air currents reaching longer distances [125,126]. If sexual reproduction occurs, even sporadically in this species, it can contribute to long distance dissemination, leading to a broader pathogen distribution.

The North American clade encompassed isolates from the United States, Canada, and one strain from Africa obtained from a culture collection. This cluster is more genetically diverse and seems to have a separate evolutionary history. Although the first report in the literature of a *Colletotrichum* species infecting maize came from Europe (Italy, 1852) we argue that the North American cluster may be the more ancient lineage (or ancestral lineage) since maize anthracnose was described for the first time in 1914 in the United States [127]. Considering that Mesoamerica is the maize center of origin, most probably Mexico, [128,129] and the center of origin of pathogenic species usually correspond to the center of origin of their host [130], given the proximity to the USA, this lineage could be the founder of the outbreak during the 1970s and may have served as the source for introductions in other continents. The highest genetic diversity found in this lineage also corroborates this hypothesis, since the center of origin of a pathogen is likely to consist of populations with more genetic variability than recently founded populations [131]. *C. graminicola* may have evolved together with the host in Central America, and due to the expansion of maize production in the USA green belt the pathogen had an opportunity to spread across the vast maize growing areas. Alternatively, our sampling, which consisted of a large range of years (1975 to 2016), might have overestimated the genetic diversity of this group. A more contemporary sampling can reveal the current genetic variation of populations from North America.

In South America, isolates from Argentina and Brazil were grouped into separate clades, suggesting independent introductions. In Brazil, the first disease report was in 1965 in the São Paulo state (Campina’s city) [132]. In the country, maize is planted in two distinct growing seasons, the first crop planted in the summer and the second crop during the winter. This practice contributes to increasing primary inoculum in the area. A significant reduction in maize yield in tested hybrids was reported by Cota et al. [43] which was associated, among other factors, with a higher pathogen aggressiveness and genetic differentiation in populations during the crop cycle. This could indicate changes in the population size and their genetic composition that may have contributed to a higher disease virulence in the country.

We found molecular evidence of genetic recombination for the three *C. graminicola* populations detected in this study. The rapid decay of linkage disequilibrium and extensive reticulations in the neighbor network is consistent with a possible history of recombination. Furthermore, the PHI test for clonality detected recombination within each population (*P* = 0.0). These results corroborate a previous study that found evidence of recombination using the PHI test at the chromosome level [92]. Similar findings of low and fast LDecay are reported in other fungal plant pathogens which suggests the occurrence of genetic mechanisms responsible for creating such patterns in genomes [82,133,134]. In addition, the signature of repeat-induced point mutation (RIP), a genome defense mechanism that occurs during sexual reproduction, [135,136] was detected in *C. graminicola* genomes [137,138]. RIP mutations may indicate an ancestral state of sexual reproduction or that meiosis cryptically occurs in nature [71,139].

Despite *C. graminicola* being reported as a homothallic species, strains may exhibit homothallism and heterothallism, but this phenomenon is not yet fully understood [22]. Investigation of fertility through crosses of the strains M1.001 and M5.001 produced an offspring exclusively heterothallic, which led Vaillancourt et al. [79] to hypothesize that unbalanced heterothallism could be involved in the mating system. Our investigation of sex-related genes revealed gene losses in most isolates and in those species known to be self-fertile such as *C. salicis,* and in those known to be self-sterile and cross-fertile like *C. fiorinae* [120]. Interestingly the only two genes missing in all *C. graminicola* genomes analyzed were *ppg1* and *pre2,* which form a pheromone receptor pair. The importance of this signaling pair varies among homothallic ascomycetes. However, none of the deletion mutants obtained in *Sordaria macrospora*, *Aspergillus nidulans* and *Gibberella zeae* completely lost the capability to sexually cross [115].

Considering this evidence and the high completeness of the assemblies we conclude that at least three of the *C. graminicola* isolates maintained all the genes required to undergo sexual reproduction. This finding also suggests that some lineages have lost the ability for sexual reproduction or that not all sex-related genes reported in other species are required in *Colletotrichum* spp. [22]. Keeping in mind the presence of homothallic and heterothallic strains in this species, the lack of genes required to sexually cross in one of the strains could mean failure in sexual reproduction. In this scenario, sexual reproduction would become rare (but not inconceivable) due to the low chance of two strains having the full set of genes encounter themselves and mating.

The reproductive biology of the genus *Colletotrichum* is mixed, and the sexual state of several species is unknown although several species are known to have the capacity for sexual reproduction. The sexual capability of *C. graminicola* is well-documented [78,140]. We demonstrated experimentally the sexual capability of *C. graminicola* isolates mating in lab conditions, as described by Vaillancourt et al. [79] (see Supplementary Fig. S4 for details), although the teleomorph has not yet been found in field conditions. Although the absence of the sexual state in nature could cause less recombination, population genetic studies show that several *Colletotrichum* species known as exclusively asexual, exhibit evidence of genetic recombination [83,141,142]. Some studies have described alternative mechanisms, such as the parasexual cycle as responsible for creating genetic recombination in *Colletotrichum* species [79,84,143–145].

Species with the potential for genetic exchange are more likely to overcome control measures employed since recombination can result in new combinations of alleles that increase diversity and speed up adaptation at the field scale [63,146]. This study gathered evidence that *C. graminicola* populations may have been affected by genetic recombination, but we also showed evidence that even if rare (possibly due to frequent losses of sex and meiosis-associated genes) *C. graminicola* can undergo sexual recombination. Fine-scale investigations at the genome level can be employed to understand the real impact of genetic recombination on genetic diversity and adaptation, for instance, if genetic exchange enhances virulence factors (also called fitness-related traits) in these populations, as suggested for other fungal species [147–151]. That can be particularly important in the agricultural pathosystem given that crop pathogens are constantly facing a homogeneous environment that imposes strong directional selection that leads to the evolution of more virulent and drug-resistant pathogen populations [130,152].

We did not find a correlation between virulence and population structure; however, we observed a substantial difference in the spectrum of virulence. Several isolates were more aggressive than the strain M1.001 which was originally collected from North America in 1972 during the US. outbreak [153]. This finding is particularly important because this strain represents the genetic diversity existent in the outbreak and strains more aggressive are currently present in nature, which may be a risk to the development of new, even more, destructive outbreaks. The pathogenicity assay was performed in one maize line (Mo940) notably susceptible to anthracnose. The use of distinct maize genotypes can result in different responses to fungal infection, as already reported in previous studies, and it should be considered in further pathogenic characterizations [42,154,155]. Another significant aspect is that resistance to maize anthracnose is inherited quantitatively, i.e., host resistance is controlled by several genes with additive genetic effects [156–158]. This fact may correlate with the range of virulence found in this study. The three genetic clades of *C. graminicola* detected may have distinct evolutionary origins, hence encompassing a diverse combination of virulence genes, which could cause different defense responses from the host. We also highlighted that the large variability in virulence should be considered for the screening of resistant maize varieties. Population studies can support breeding programs by investigations of intraspecific variability of *C. graminicola* populations since the effectiveness of genetic resistance depends on knowledge of the genetic structure of pathogens.

## Conclusions

Our study represents the first investigation of the population structure of *C. graminicola* infecting maize. The three main clades (North America, Brazil, and Europe) have different levels of genetic diversity suggesting that each has a unique evolutionary history. We also found evidence of recent migration of isolates between Europe and Argentina, possibly due to the importation of infected plant material. Although the sexual state of *C. graminicola* has only been reported in lab conditions, we show strong evidence that genetic recombination, whether by sexual reproduction, or by alternative mechanisms (such as the parasexual cycle) plays an important role in *C. graminicola* population structure. In contrast to the traditional view of *C. graminicola* being mainly asexual, we conclude that genetic recombination is frequent. Fine-scale investigations at the genome level can be employed to understand the real impact of recombination and gene flow on the genetic diversity and adaptation of this species, with a direct effect on control measures and breeding programs aiming at resistance. Finally, we warn of the danger that maize anthracnose has the potential to become much more significant in the future, mainly due to highly aggressive isolates currently present in maize fields and the geographic range expansion of the pathogen, in addition to increased susceptibility of agroecosystems accentuated by ongoing climate changes.

## Additional Files

**Supplementary Fig. S1.** A) Bayesian information criteria (BIC) indicating the most probable number of genetic groups by discriminant analysis of principal components analysis (DAPC). B) Scatterplot from DAPC of *Colletotrichum graminicola* isolates assigned with *a priori* geographical information (by continent).

**Supplementary Fig. S2.** Bayesian information criteria (BIC) indicating the most probable number of genetic groups using the SNAPCLUST function.

**Supplementary Fig. S3.** Colony morphology of *Colletotrichum graminicola* showing phenotypic diversity among isolates. The colonies were cultivated on PDA medium, at 23°C under continuous light for 6 days.

**Supplementary Fig. S4.** Sexual structures of *Colletotrichum graminicola* produced by crossing from strains M1.001 (CB130836) and M5.001 (CBS 130839). The cross-inoculation experiment was performed under laboratory conditions as described by Vaillancourt and Hanau 1991. (a) Agar plugs containing strains growing on autoclaved leaves of maize. (b) Mature perithecia, picture taken with Olympus SZX9 stereomicroscope outfitted with an Olympus DP70 camera. c) Ascus and ascospores. d) Ascospores.

**Supplementary Table S1.** Isolates of *Colletotrichum graminicola* used in this study.

**Supplementary Table S2.** Summary statistics of *Colletotrichum graminicola* genomes generated in this study.

**Supplementary Table S3.** Measure of pairwise genetic differentiation between genetic clusters of *Colletotrichum graminicola*.

**Supplementary Table S4.** Multilocus genotypes (MLGs) among *Colletotrichum graminicola* isolates.

**Supplementary Table S5**. *Colletotrichum graminicola* orthologs of *Neurospora crassa, Saccharomyces cerevisiae, Podospora anserina, Fusarium graminearum*, and *Schizosaccharomyces pombe* sex-related genes.

## Data Availability

The vcf file generated in this study is provided as Additional file 1. The genome assemblies were submitted to GenBank, and their accession numbers are in Supplementary Table S2.

## Abbreviations

ANOVA: Analysis of Variance
BIC: Bayesian information criterion
BLAST: Basic Local Alignment Search Tool
BR: Brazilian clade
CTAB: cetyltrimethylammonium-bromide
EU: European clade
FST: Genetic differentiation index
GTR: general Time-Reversible model of nucleotide substitution
ITS: internal transcribed spacer
MAF: minor allele frequency
MLGs: multilocus genotypes
NAM: North American clade
MSN: minimum spanning network
NCBI: National Center for Biotechnology Information
PDA: potato dextrose agar
PDB: potato dextrose broth
PCA: principal component analysis
PHI: pairwise homoplasy index
RAD-Seq: restriction site-associated DNA sequencing; pairwise homoplasy index
SNPs: single-nucleotide polymorphisms
WGS: whole-genome sequencing

## Authors’ Contributions

S.A.S. and M.R.T conceived the study and designed the project; S.A.S performed fungal isolation and DNA extraction; M.R.T., F.R., R.B., S.B. J.L.V.V., S.A.S, and F.B.C.F. performed analyses; M.R.T., F.R., R.B, and S.A.S. wrote the manuscript; F.B.C.F. and S.A.S. performed the pathogenic characterization; M.A.A.P., M.M.W. and J.A.C. performed genome sequencing; I.B., W.B., V.O., J.B., A.T., J.S.D., P.D.E., P.R., T.J.Z., J.H., G.M. collected the samples. All authors reviewed the manuscript.

## Acknowledgments

The authors thank Maria Elena Bayón Sánchez for her assistance in the isolation of samples and pathogenicity tests, Lisa J. Vaillancourt, and the Genetic Resources Center (NARO) for providing some isolates.

