## Supplementary Fig. S1; Supplementary Fig. S2; Supplementary Fig. S3; Supplementary Fig. S4 for "Population genomics provide insights into the global genetic structure of *Colletotrichum graminicola,* the causal agent of maize anthracnose"

[1]Instituto de Investigación en Agrobiotecnología (CIALE), Departamento de Microbiología y Genética, Universidad de Salamanca, Villamayor, 37185 Salamanca, Spain. [2] Department of Agricultural and Food Sciences (DISTAL), University of Bologna, Viale Fanin 44, 40126 Bologna, Italy. [3]United States Department of Agriculture, Foreign Disease and Weed Science Unit, 1301 Ditto Avenue, Fort Detrick MD, 21702 USA. [4] Embrapa Environment, Jaguariúna, SP, Brazil. [5] Center for Gastrointestinal Biology and Disease, Division of Gastroenterology and Hepatology, and UNC Microbiome Core, Department of Medicine, School of Medicine, University of North Carolina, Chapel Hill, NC 27599, USA. [6] USDA-Animal and Plant Health Inspection Services, Biotechnology Regulatory Services, Riverdale, MD 20737, USA. [7]Syngenta Seeds La Grangette, 32220 Lombez France. [8] Bayer Crop Science/Monsanto SAS 82170, Monbequi, France. [9] Ontario Ministry of Agriculture, Food, and Rural Affairs, University of Guelph-Ridgetown, Ridgetown, Ontario, Canada. [10] Facultad de Ciencias Exactas Físicas y Naturales, Universidad Nacional de Córdoba, IMBIV-CONICET-ICTA Av Vélez Sarsfield 1611, Ciudad Universitaria, Córdoba, Argentina. [11] Department of Plant Pathology and Environmental Microbiology, Pennsylvania State University, State College, PA, 16801, United States. [12] Misión Biológica de Galicia, Spanish National Research Council (CSIC), PO Box 2836080, Pontevedra, Spain. [13] Department of Plant Pathology, University of Nebraska–Lincoln, 68583-0722. [14] Federal Department of Economic Affairs, Agroscope Reckenholzstrasse 191, 8046 Zurich, Switzerland. [15] Department of Plant Pathology and Microbiology, Iowa State University, Ames, IA 5001. [16] Bc Institute for Breeding and Production of Field Crops, 10370 Dugo Selo, Croatia. [17] Statistics Department University of Salamanca, Salamanca, Spain.

***Corresponding authors:** Serenella A. Sukno, Institute for Agribiotechnology Research (CIALE), Departamento de Microbiología y Genética, Universidad de Salamanca, Villamayor, 37185 Salamanca, Spain. https://orcid.org/0000-0003-3248-6490; Michael R. Thon, Institute for Agribiotechnology Research (CIALE), Departamento de Microbiología y Genética, Universidad de Salamanca, Villamayor, 37185 Salamanca, Spain. https://orcid.org/0000-0002-7225-7003

Table of contents:


| | |
|---|---|
| Figure S1 | Page 2 |
| Figure S2 | Page 3 |
| Figure S3 | Page 4 |
| Figure S4 | Page 5 |

A)

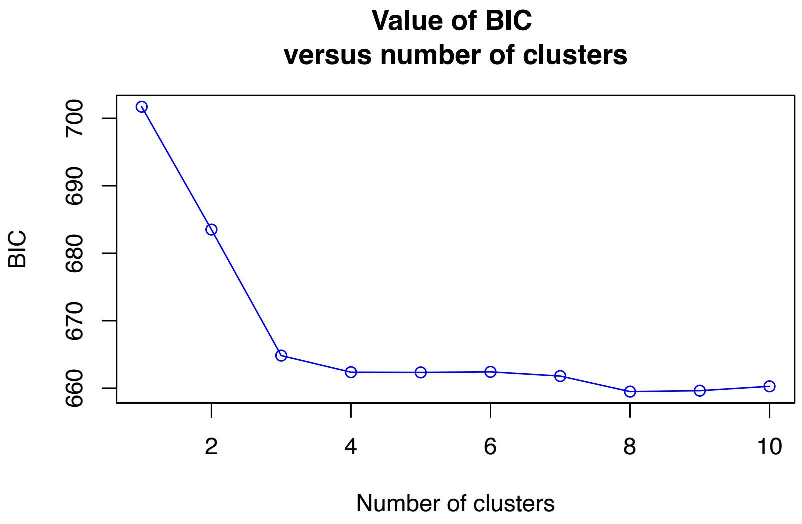


B)

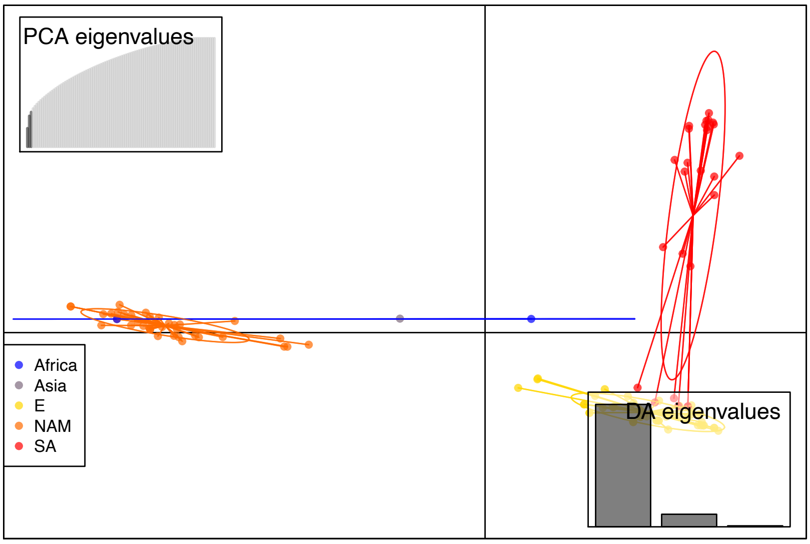


**Supplementary Fig. S1.** A) Bayesian information criteria (BIC) indicating the most probable number of genetic groups by discriminant analysis of principal components analysis (DAPC). B) Scatterplot from DAPC of *Colletotrichum graminicola* isolates assigned with a priori geographical information (by continent).

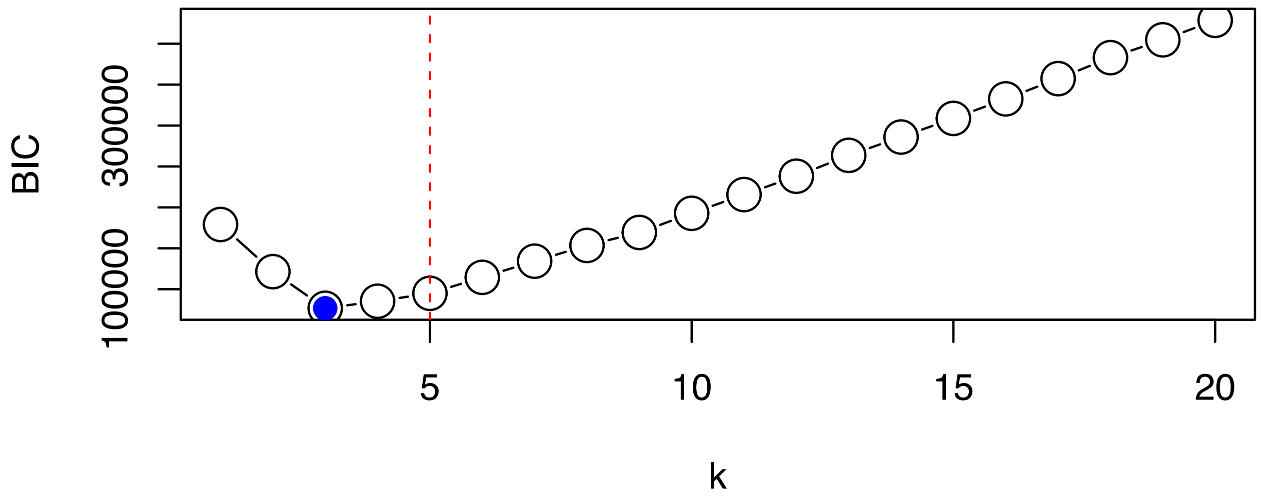


**Supplementary Fig. S2.** Bayesian information criteria (BIC) indicating the most probable number of genetic groups by SNAPCLUST function.

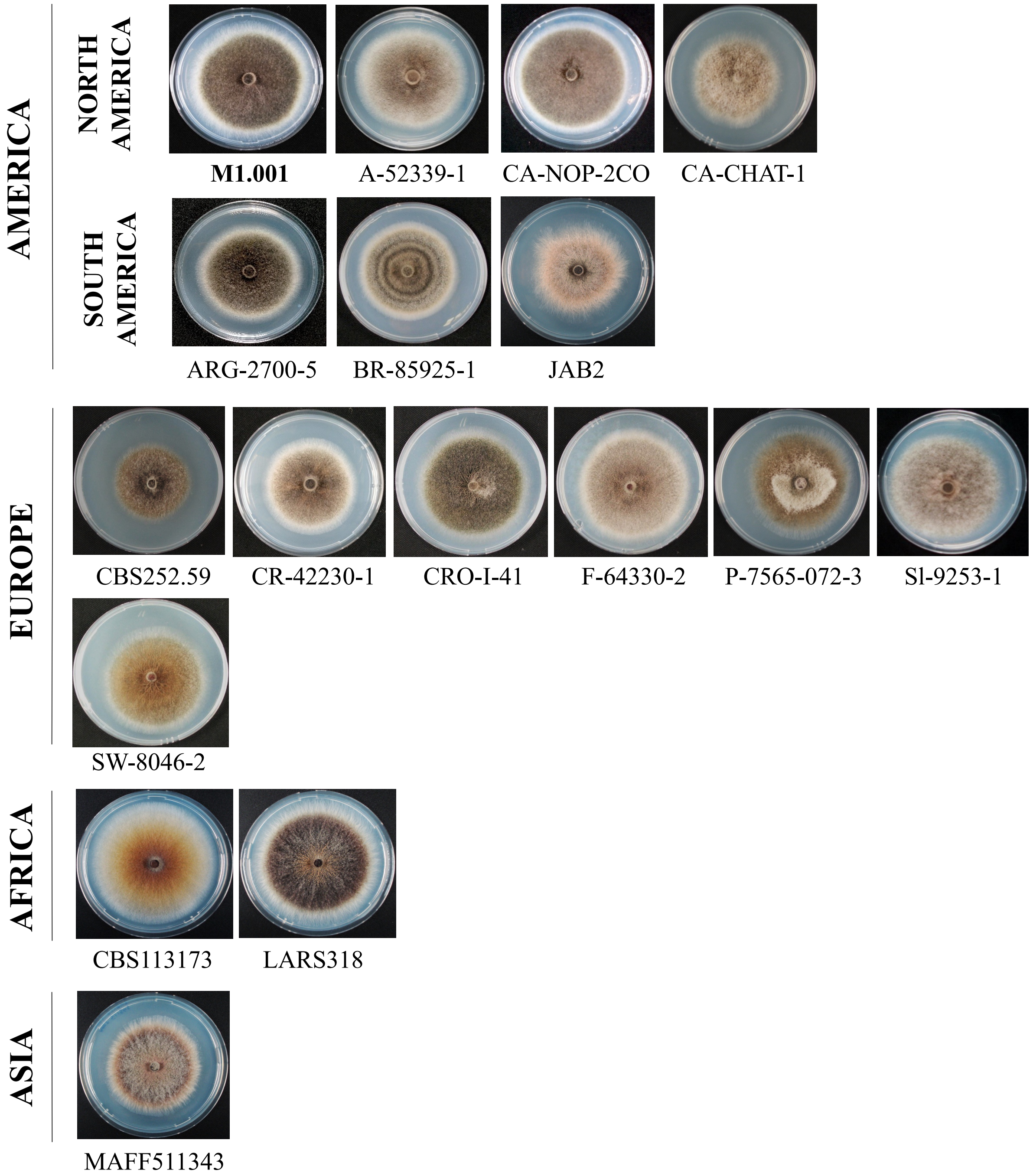


**Supplementary Fig. S3.** Colony morphology of *Colletotrichum graminicola* showing phenotypic diversity among isolates. The colonies were cultivated on PDA medium, at 23ºC under continuous light for 6 days.

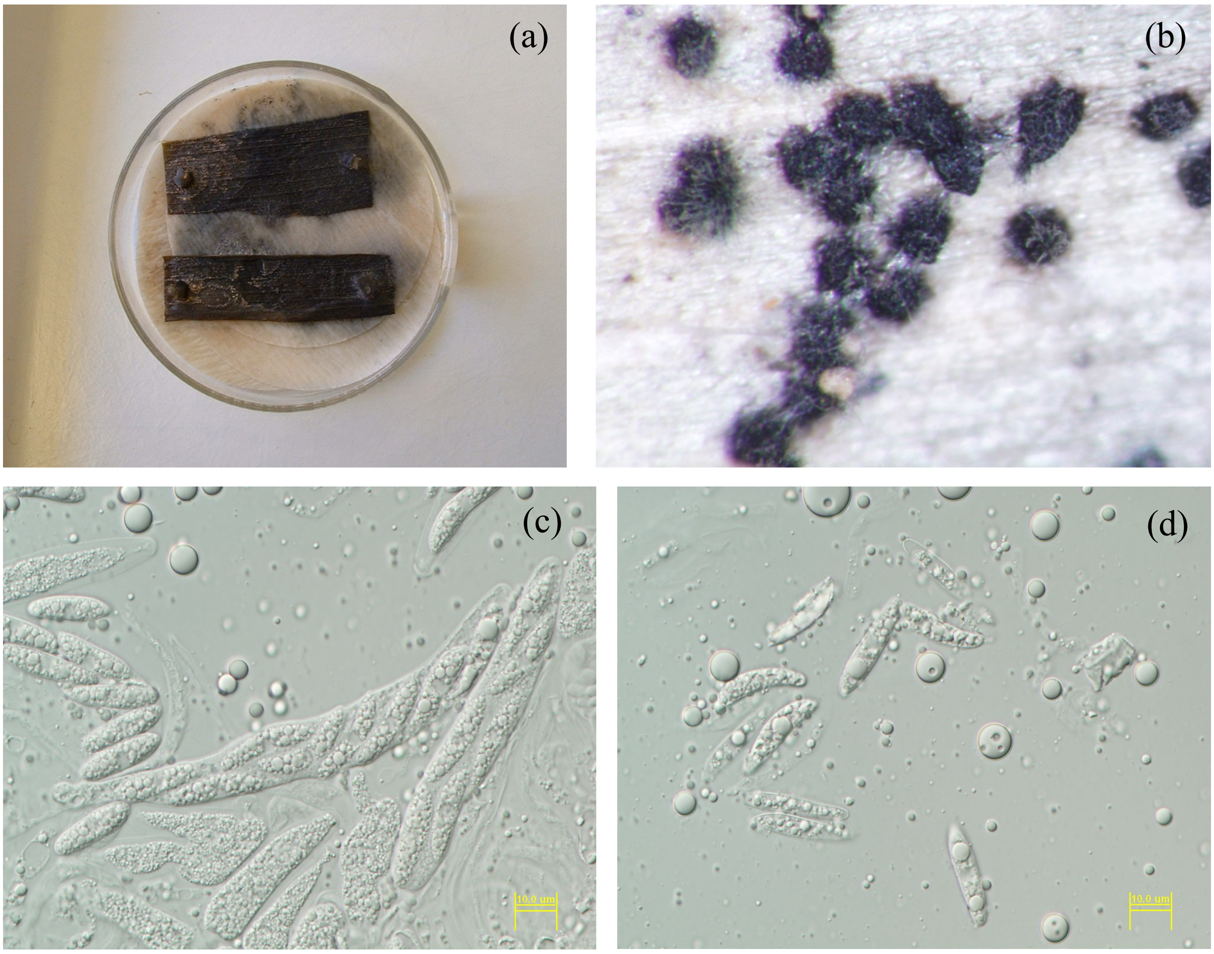


**Supplementary Fig. S4.** Sexual structures of *Colletotrichum graminicola* produced by crossing from strains M1.001(CB130836) and M5.001 (CBS 130839). The cross-inoculation experiment was performed under laboratory conditions as described by Vaillancourt and Hanau 1991. (a) Agar plugs containing strains growing on autoclaved leaves of maize. (b) Mature perithecia, picture taken with Olympus SZX9 stereomicroscope outfitted with an Olympus DP70 camera. c) Ascus and ascospores. d) Ascospores
